# Visual Search P300 Source Analysis Based On ERP-fMRI Integration

**DOI:** 10.1101/2020.07.16.206375

**Authors:** Qiuzhu Zhang, Cimei Luo, Junjun Zhang, Zhenlan Jin, Ling Li

**Author notes:** Corresponding author: Ling Li.

## Abstract

Attention control can be achieved in two ways, stimulus-driven bottom-up attention and goal-driven top-down attention. Different visual search tasks involve different attention control. The pop-out task requires more bottom-up attention, whereas the search task involves more top-down attention. P300 which is the positive potential generated by the brain in the latency of 300-600 ms after the stimulus, reflects the processing of cognitive process and is an important component in visual attention. The P300 source is not consistent in the previous researches, our aim therefore, is to study the source location of P300 component based on visual search attention process. Here we use pop-out and search paradigm to get the ERP data of 13 subjects and the fMRI data of 25 subjects, and analyze the source location of P300 using the ERP-fMRI integration technology with high temporal resolution and high spatial resolution. The target differs from the distractor in color and orientation in the pop-out task, whereas the target and the distractor have different orientation and the same color in the search task. ERP results indicate that pop-out induces larger P300 concentrated in the parietal lobe, whereas search induced P300 is more distributed in the frontal lobe. Further ERP and fMRI integration analyses reveal that the left angular gyrus, right postcentral gyrus of parietal lobe and the left superior frontal gyrus (medial orbital) are the source of P300. Our study suggests the contribution of the frontal and parietal lobes to the P300 component.

## I. INTRODUCTION

Attention plays very important role in our daily life. For example, when driving, the driver pays attention to the road conditions while the teacher pays attention to the student performance during the lesson, and the doctor pays attention to the patients’ symptoms and write prescription during treatment. Precisely, attention entails directing concentration selectively towards certain stimulus while ignoring others, and it is a common psychological feature accompanied by psychological processes such as sensory perception, memory, thinking, and imagination. According to the biased competition model, there are two ways of attention control. The first one is bottom-up attention which is driven by stimulus, also known as exogenous attention and the second one is top-down attention which is driven by a goal, also known as endogenous attention [1, 2]. Top-down attention needs to be motivated and maintained by individuals on purpose, so it takes a certain amount of time and cognitive effort. On the other hand, bottom-up attention reflects the mechanism of the salient stimulus’s automatic attention bias, so it is automatic, fast and effortless [3].

Furthermore, visual attention guides visual search. In essence, feature integration theory, guided search theory, and similarity theory provide the theoretical basis for visual search. The feature integration theory contains two basic concepts, object and feature. The feature refers to a specific value of a dimension (such as color, orientation, and shape), whereas the object refers to the combination of some features (such as the red square forming the object). Visual process is divided into two stages including pre-attention (i.e., feature registration) and feature integration (i.e., object of consciousness). The proponents of this theory argue that features are automatically processed in parallel in the pre-attention stage, while the features are integrated into objects in a series of top-down processing in the integration stage [4]. On the basis of the feature integration theory, researchers successively proposed the guided search theory [5] and the similarity theory [6]. According to the guided search theory, the sequence of the second stage series processing depends on the results of the first stage parallel processing, and the series processing is carried out under the guidance of the results of parallel processing. For the similarity theory, the difficulty of visual search depends on the similarity of stimulus materials, and the lower the similarity between the target and the distractor, the easier it is to search for the target.

Attention mechanism can be studied electro-physiologically using event-related potentials (ERP). ERP is a basic, non-invasive electrophysiological detection method, and it plays an increasingly important role in understanding the mechanism of attention [7, 8]. Specifically, ERP components include exogenous (physiological) and endogenous (psychological) components. P300 is the most involved endogenous component, which is not affected by the physical characteristics of the stimulus and is related to the mental state and attention of the subjects. To be precise, P300 is the positive potential generated by the brain in the latency of 300-600 ms after the presentation of stimulus, which reflects the processing of cognitive process [9]. Essentially, P300 can be divided into P3a and P3b, where P3a is an assessment of the initial state of a stimulus, recording the time it takes for the stimulus to appear until the stimulus awakenings, whereas P3b is the time it takes for the signal to attract the attention and memory [10, 11]. Despite the relatively large number of studies on the source of P300, the source location is still not consistent. The P300 component was initially proposed to be caused by the performance of individual target stimulus in the oddball paradigm, and showed its maximum positive peak at 300 ms after the target was triggered, with obvious activation in the parietal cortex [9]. By using the classic oddball paradigm to induce P300, it was found that the activated brain regions included the anterior cingulate, middle frontal gyrus, angular gyrus, middle temporal gyrus, hippocampus/parahippocampal gyrus, and temporal parietal junction [12, 13]. The early source studies obtained through ERP were uncertain, researchers used functional magnetic resonance imaging (fMRI) to study the P300 source and found that both auditory and visual stimuli involve the bilateral superior gyrus and anterior cingulate gyrus, two brain regions with relatively consistent activation [14].

Since P300 contains P3a and P3b, recent studies have used low resolution electromagnetic tomography (LORETA) to analyze the generation of P300 components and found that P3a and P3b showed different scalp morphology and cortical sources [15, 16]. In terms of scalp morphology, the scalp distribution position of P3a is more anterior than that of P3b. Whereas in terms of cortical source, the source of P3a is located in the cingulate, frontal and right parietal lobes, whereas the source of P3b includes bilateral frontal lobes, parietal lobes, limbic region, cingulate and temporal-occipital regions [15-17]. The P300 component induced by auditory materials also show differences in source location. The source location of P3a include the anterior cingulate, frontal and parietal lobes, whereas the source location of P3b is distributed in a wider network, including the superior temporal gyrus, middle temporal gyrus, posterior parietal cortex, hippocampus, cingulate and frontal lobes [18, 19]. Although the current source study on P300 focuses on healthy people, clinical studies are also on the rise. A study on patients with schizophrenia found that P3a was more distributed in the frontal lobe, and P3b was less activated in the frontal lobe and cingulate cortex [16]. Another study used standardized low resolution electromagnetic tomography (sLORETA) to compare current source density (CSD) in patients with post-traumatic stress disorder (PTSD) and healthy controls, and found that P300 current source density in PTSD patients was significantly reduced in the inferior frontal gyrus, precentral gyrus, insula, and anterior cingulate [20].

ERP and fMRI are commonly used techniques to measure changes in neural activity of brain function. Electroencephalogram (EEG) signals are instantaneous, with millisecond resolution. There is little delay effect on neural events, but the spatial resolution is poor due to the role of volume conductors such as scalp, skull and cerebrospinal fluid [21]. Conversely, the function magnetic resonance imaging (fMRI) measures the signal of blood oxygenation level dependent (BOLD) in the process of brain activity. But it generally takes several seconds time from neural activity to hemodynamic response, resulting in poor temporal resolution. However, the use of integrated ERP-fMRI enables the harmonization of the recorded EEG with the BOLD signal to achieve high temporal resolution and high spatial resolution [22-24]. ERP-fMRI is considered to be an effective cognitive neural technique for studying ERP components [15, 25]. Actually, it is possible to use ERP-fMRI integration technology to study neural activity more accurately.

In the present study, we use pop-out and search visual search task to study attention-related ERP component. We perform ERP and fMRI experiment based on visual search task separately, and use ERP-fMRI integration technology to analyze P300 source. Activation regions of fMRI are mapped to the ERP difference source model, and the waveform and topographic map of the corresponding source regions are extracted to determine the brain region producing P300. We assumed that the pop-out and search conditions induced P300, but P300 of the two conditions was different. The activity of the frontal-parietal lobe was related to the source of P300.

## II. METHODS

### A. ERP EXPERIMENT BASED ON VISUAL SEARCH

#### 1) Participants

Fourteen healthy subjects (7 females, mean age = 24 years; age range 18-30 years) were recruited to participate in the study. All subjects were right-handed, with normal vision or corrected to normal vision, and had no history of neurological disease. Since one female was excluded from analyses because of excessive blinks (more than 3% trials), valid data were analyzed from the 13 remaining subjects. The research was approved by the local human research ethics committee. All subjects gave written informed consent. In addition, all subjects were informed of the detailed requirements before they began the experiment. At the end of the experiment, all subjects were given monetary compensation.

#### 2) Stimulus and procedure

The study was within-subjects design, and all subjects were required to complete the 2 conditions (pop-out and search conditions). In the pop-out condition, the target triangle was different from the other three distractor triangles in color and orientation. In the search condition, the color of the target triangle was the same as other three distractor triangles, but the orientation was different. The stimulus materials were 16 isosceles triangles with 8 different orientations and 2 colors (red and green). The triangle got eight different orientations by rotating it at different angles (clockwise 0°, 45°, 90°, 135°, 180°, 225°, 270°, 315°). Additionally, each triangle had the same area, brightness and saturation.

Figure 1 illustrates an example of the two conditions. A fixation cross of 500 ms was presented, and then the target triangle (sample) 1000 ms. The target triangle only appeared once in each session. Subjects were instructed to maintain attention and remember the color and orientation of the target triangle for the judgment. Following the target triangle (sample), there was a 500 ms delay and then the presentation of the visual array with four stimuli composed of three distractors and the target. In this process, subjects were required to make key response. When the target triangle appeared on the left side of fixation cross, they were required to press “1” key, and when the target triangle appeared on the right side of fixation cross, to press “2” key. During the whole process, the subjects had to stay focused and use the peripheral light to focus on the left and right field of vision in the center of the screen to make quick and accurate judgments. The number of trials and target locations of pop-out and search conditions were balanced in the experiment. Subjects took formal trials after completing 64 training trials, which consisted of 12 blocks, each containing 32 trials, for a total of 384 trials. Each block took about 2.5 minutes, and the interval time of each block was 1min. The total experimental time was about 40 minutes. The stimuli were presented using E-prime1.1 software.

**FIGURE 1.**
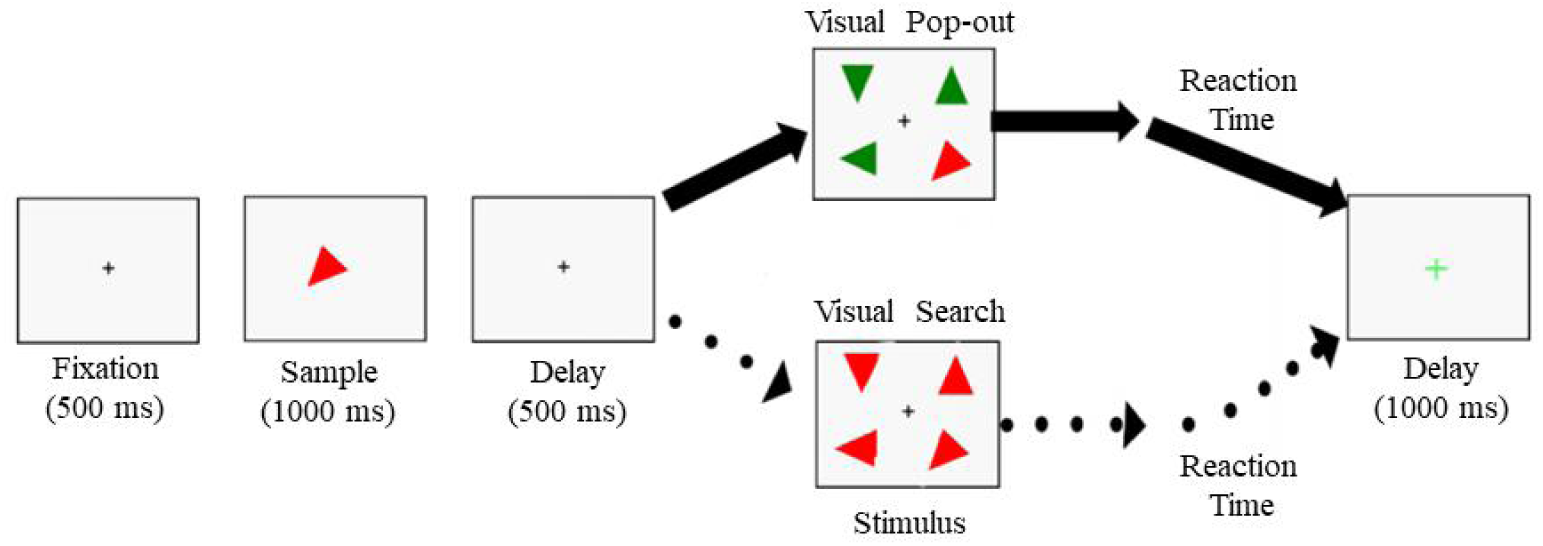
Visual search paradigm. In the pop-out condition, the target triangle was different from the other three distractor triangles in color and orientation. In the search condition, the color of the target triangle was the same as other three distractor triangles, but the orientation was different. Each trial began with a fixation cross, and then the target triangle (sample) was presented. Following the target triangle (sample), there was a delay and then the presentation of the visual array with four triangles composed of three distractors and the target. Subjects were to make a judgment whether the target triangle appears on the left or right side of the fixation cross. After reaction, a delay was presented.

#### 3) ERP data recording and analysis

ERP data were recorded using 64 channels ActiveTwo system (Biosemi, The Netherlands). All channels were amplified with an analog bandpass filter of 0.06-208 Hz and were digitized at 1024 Hz. At the same time, two earlobes and four electrooculogram (EOG) channels were recorded.

ERP data were processed and analyzed using MATLAB software. The two-way FIR bandpass filter of 0.5-55 Hz in EEGLAB toolbox was used to filter the EEG signal, and the EEG signal was segmented from 200 ms before the stimulus onset to 1000 ms after the stimulus onset. We removed the eye movement artifacts, and the amplitude whose difference between two vertical EOG and/or the two horizontal EOG was greater than 100µV, or the signal whose mean value of EOG was more than 3 times the standard deviation. In addition, trials with the wrong response and reaction time beyond 200-1200 ms were invalidated. After preprocessing, the valid trials of all subjects under each condition was more than 25. We selected 300-600 ms time window and 6 central-parietal electrodes (CP1, CPz, CP2, P1, Pz, P2) to analyze P300 component [26, 27], comparing the differences of P300 component under different conditions.

### B. fMRI EXPERIMENT BASED ON VISUAL SEARCH

#### 1) Participants

Twenty-six right-handed, healthy subjects (8 females, mean age = 21.6 years, age range 18-24 years) participated in the study. All subjects had normal or corrected to normal vision, and had no history of neurological disease. All subjects were informed about the requirements of the experiment in detail before the experiment. They were required to sign a series of surveys to eliminate all possible risks and maximize safety (such as whether the subjects had metal in their bodies). More so, subjects were scanned before entering the laboratory to prevent metal objects from being brought into the laboratory. The study was approved by the Ethics Committee of the University of Electronic Science and Technology of China. All subjects gave written informed consent. At the end of the experiment, all subjects were given monetary compensation.

#### 2) Procedure

The fMRI experiment was modified from ERP experiment. The experiment was a within-subjects design, including the pop-out and the search task. The experiment further adopted a block design, dividing the stimulus into different types while combining the same type of stimulus into a block, and there after presenting the stimulus in block form. The stimulus presented in each block was continuous and repeated. When the task stimulus is presented for a long time, the amplitude of the signal of BOLD signal is higher, therefore, the signals of multiple blocks are superimposed, thus the BOLD signal varied greatly [28].

There were 8 blocks for each pop-out task and search task, and the same task trial repeated 4 times in each block (that is, 4 trials of the same type occurred after a sample), so there were 32 trials for each of the two tasks. In order to avoid the fatigue effect of a single stimulus, subjects were presented with alternating blocks. The experimental paradigm of a single block is shown in Figure 2. The whole experimental process of fMRI is shown in Figure 3. During each block, the first sample stimulus 1500 ms matching the type appeared, and the subjects were required to remember the color and orientation of the target triangle. Then a 500 ms delay was presented, followed by a task stimulus presentation. The subjects were required to press the button and judge whether the target triangle was on the left or right side of the fixation across. When the target triangle appeared to the left of the fixation across, the subjects used the left hand to press the “1” key. When the target triangle appeared on the right side of the fixation across, they were to press the “6” key with the right hand, and the subjects were instructed to perform the same type of visual search task four times in each block. The two tasks alternated 7 times for each task. Subjects were to make fast and accurate response. The subjects performed the formal trials after completing the exercise trials. The stimulus materials were presented using E-prime 2.0 (Psychology Software Tools, Pittsburgh, USA).

**FIGURE 2.**
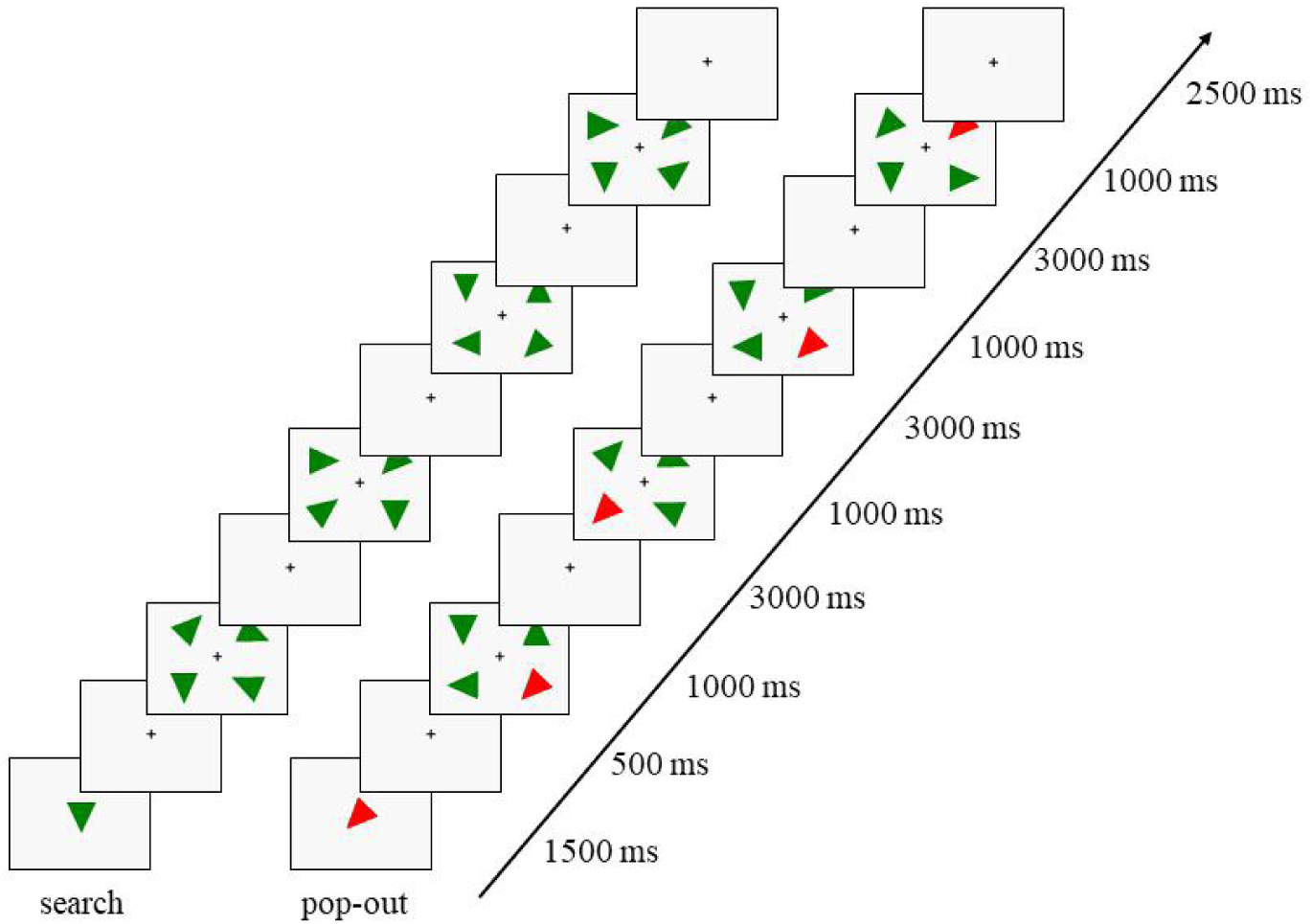
Single trial of visual search paradigm (pop-out and search tasks). The target triangle (sample) was presented. Following the target triangle (sample), there was a delay and then the presentation of the visual array with four triangles composed of three distractors and the target. The previous fixation lasted for 2500 ms, and the next trial has a delay for 500 ms, so the interval of trial was 3000 ms. Subjects were to make a judgment whether the target triangle appeared on the left or right side of the fixation.

**FIGURE 3.**
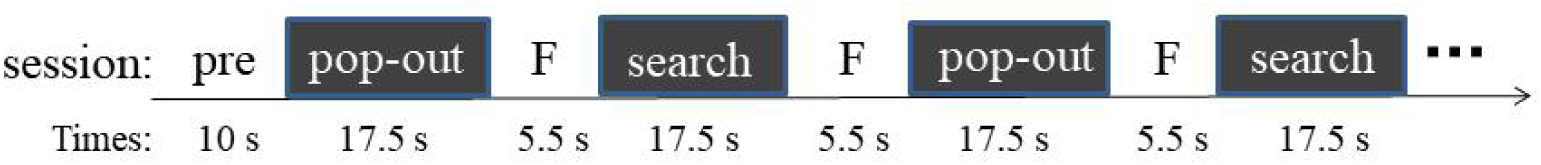
fMRI-based experimental process of visual search task. “F” represents the interval timer of each block.

#### 3) fMRI data recording and analysis

The collection of fMRI data was completed by the 3.0 T GE Discovery MR750 scanner (General Electric, Milwaukee, WI, USA), and functional images were collected by echo-planar imaging (EPI) and 8-channel phase array head coil. The corresponding parameters of fMRI image acquisition: TR= 2000 ms, TE = 30 ms, FOV= 240 × 240 mm, FA = 90°, matrix size = 64 × 64, slice thickness = 3.75 mm, slice gap = 0.6 mm, 43 slices, voxel size = 3.75 mm × 3.75 mm × 3 mm. We also collected T1-weighted images (voxel size = 1 mm × 1 mm × 1 mm, 176 slices).

For the preprocessing of fMRI data, DPARSF (Data Processing Assistant for rare-state fMRI) toolbox in MATLAB software was used [29]. In order for the subjects to adapt to the scanning environment and achieve the magnetization balance of the scanner, the first 5 time points were removed. The preprocessing included slice timing correction, realignment, spatial coregister standardization, and spatial smoothing. One subject with a total vector motion > 1.5 mm and rotation > 1.5°.

The fMRI data were analyzed using the SPM12 (statistical parametric mapping software, http://www.fil.ion.ucl.ac.uk/spm). Firstly, individual’s data were analyzed by first-level, and general linear model (GLM) was established using time related BOLD signal and regression variables. The general linear model of this experiment includes 2 regression parameters (pop-out and search) and 6 head realignment parameters (3 translational parameters and 3 rotational parameters). The group analysis was performed by second-level, and one-sample t-test of was used to obtain the activated brain areas under different conditions. False discovery rate (FDR) was used for multiple comparison correction (p < 0.05, cluster size > 25).

### C. INTEGRATION OF ERP AND fMRI

ERP and fMRI data obtained by visual search paradigm (pop-out and search task) were analyzed as follows: (1) source model reconstruction of ERP data to determine its source location; (2) significantly activated brain regions obtained by analyzing fMRI data; (3) the significant activation region obtained in the fMRI experiment was used as the source region, which was mapped to the ERP difference source model, and the waveform and topographic map of the corresponding source region were extracted to determine the brain region producing P300 component (Figure 4).

**FIGURE 4.**
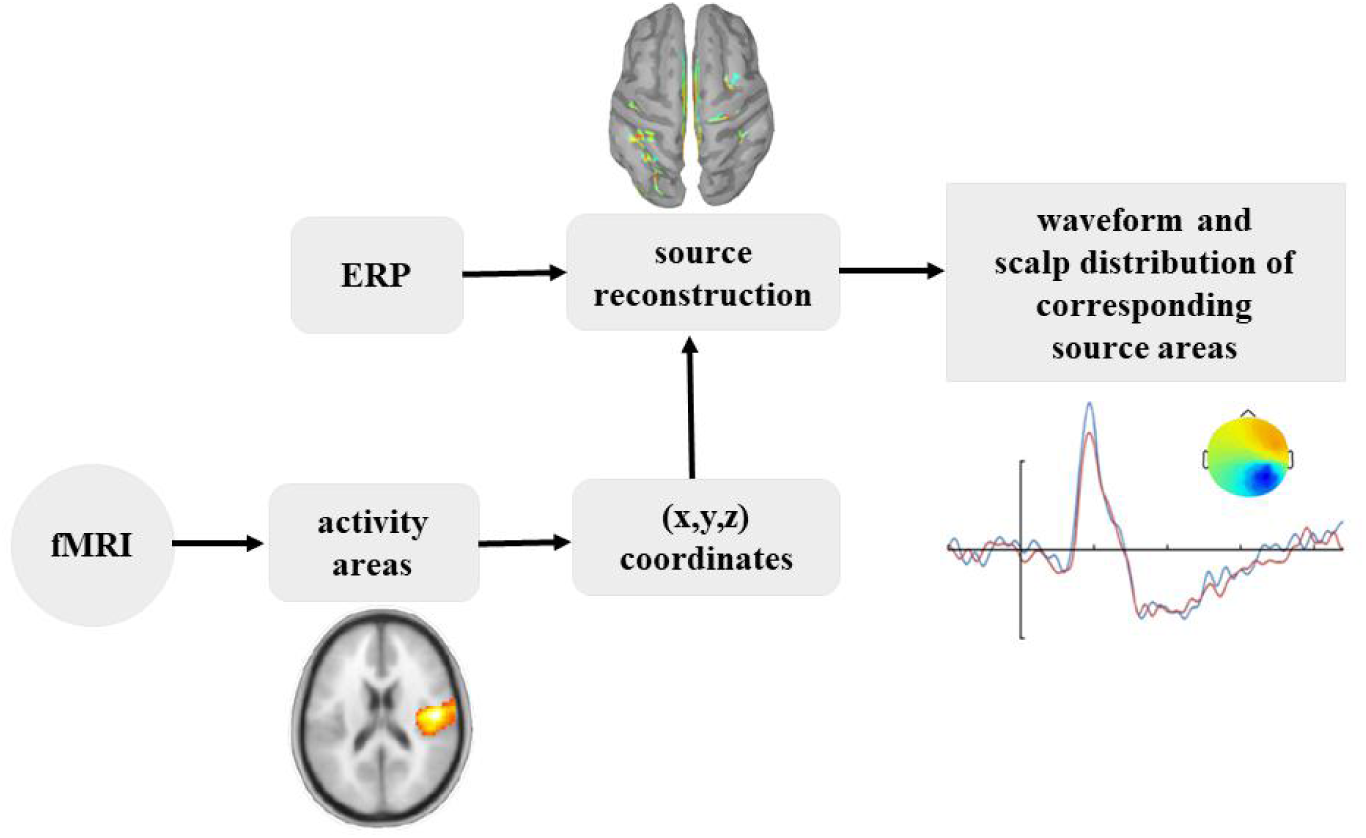
ERP-fMRI integration analysis process.

By using Brainstorm software (https://neuroimage.usc.edu/brainstorm/) source analysis was carried out on the ERP data. Selecting the Montreal Neurological Institute (MNI) template of Colin27 and using OpenMEEG to build the head model, a brain distribution source containing 15002 dipoles was established. WMNE (weighted minimum norm estimate) algorithm was used to calculate the EEG reverse problem. First, source was calculated for each subject under two different conditions, and z-value was calculated by selecting -200 to 1000 ms as the baseline. Then, the source corresponding to the same task condition of all the subjects were analyzed on the group average to obtain a group-level source activation diagram. Finally, Gaussian smoothing was applied to the group level activation graph. Since this study explored P300, we focused on the source activation diagram of the differential source model at 300-600 ms. By analyzing the ERP data of visual search, we chose the difference source model of time window of 300-400 ms for average analysis.

## III. RESULTS

### A. ERP RESULTS BASED ON VISUAL SEARCH

By analyzing the ERP signals generated by the central-parietal electrodes (CP1, CPz, CP2, P1, Pz, P2), it was found that both pop-out and search conditions induced obvious positive component within 300-600 ms time window (Figure 5). The P300 peak amplitude of 6 electrodes under pop-out condition (CP1: 12.03 ± 4.36 µV, CPz: 13.52 ± 4.8 µV, CP2: 13.89 ± 4.06 µV, P1: 13.57 ± 4.79 µV, Pz: 13.67 ± 5.6 µV, P2: 14.13 ± 4.34 µV) was higher than search condition (CP1: 7.02 ± 3.06 µV, CPz: 7.48 ± 3.39 µV, CP2: 6.84 ± 3.13 µV, P1: 6.55 ± 3.1µV, Pz: 7.09 ± 3.57 µV, P2: 6.63 ± 3.25 µV). The paired t-test established that the P300 peak amplitude in pop-out condition was significantly higher than the search condition (p < 0.001).

**FIGURE 5.**
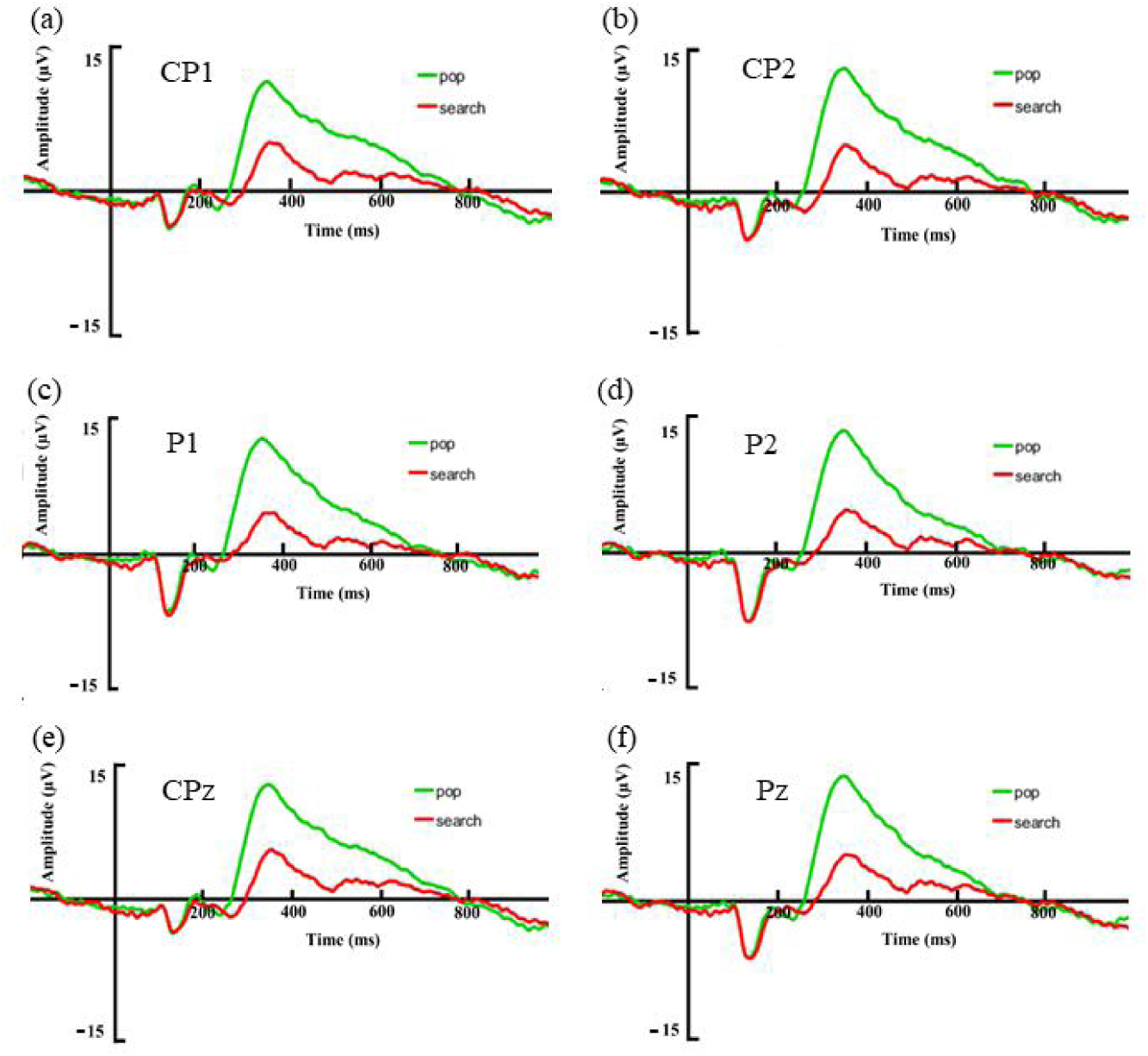
Grand average event-related potentials (ERPs) elicited by pop-out and search conditions in the 13 subjects at six central-parietal electrodes (CP1、CP2、P1、P2、CPz、Pz). Green represents pop-out condition and red represents search condition.

In addition, the P300 peak amplitude in each electrode is about 350 ms, and the range of peak time window was concentrated at 300-400 ms (350 ± 50 ms). Therefore, we narrowed the P300 time window to the range of peak time window for study.

According to the brain topographic map obtained by electroencephalogram analysis of subjects under pop-out and search conditions, it was found that P300 component in pop-out condition were mainly distributed in the parietal lobe, whereas P300 components in search condition were more distributed in the frontal lobe (Figure 6). The mean peak amplitude of P300 at the central-parietal electrodes (CP1, CPz, CP2, P1, Pz, P2) was different in two visual search tasks. The mean peak amplitude of P300 under the pop-out condition was significantly higher than that under the search condition (pop-out: 13.47 ± 4.43 µV, search: 6.94 ± 3.06 µV, t (12) = 6.598, p < 0.001).

**FIGURE 6.**
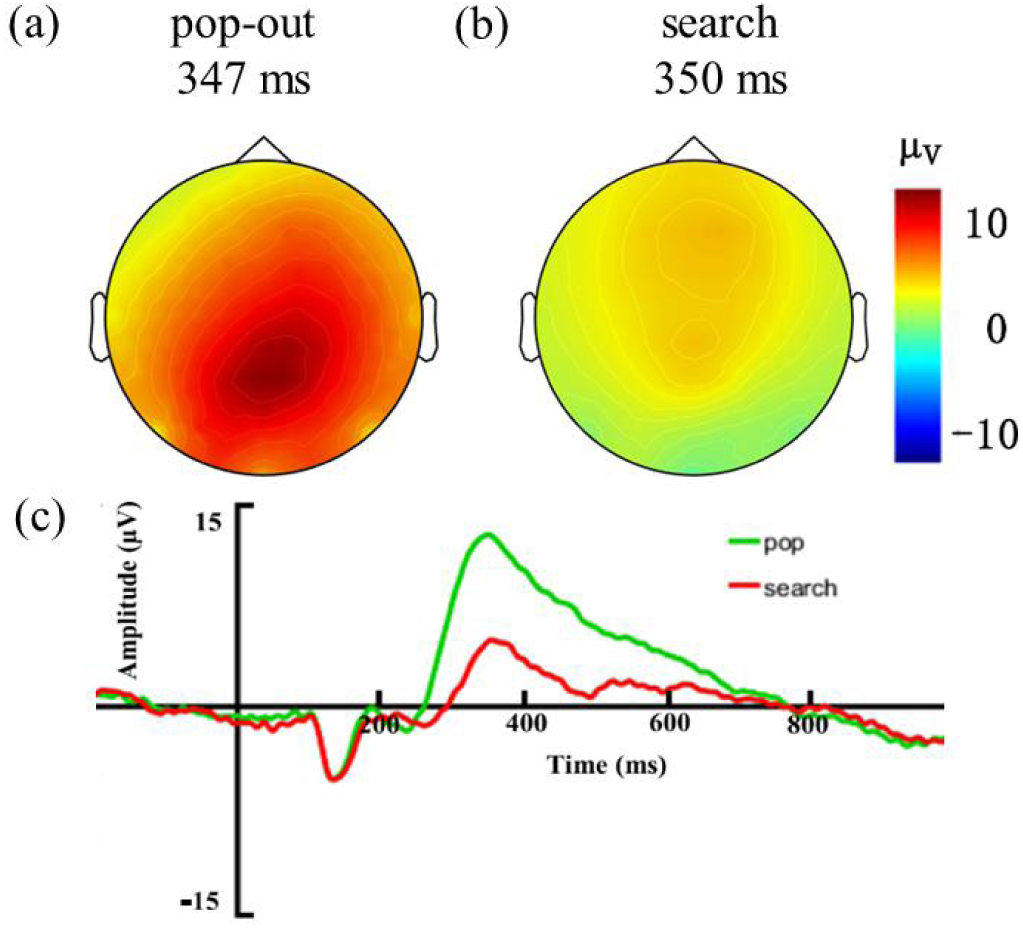
(a) Brain topographic map of P300 component (at 347 ms) of the pop-out condition. (b) Brain topographic map of P300 component (at 350 ms) of the search condition. (c) Grand average event-related potentials (ERPs) of central-parietal electrodes under the pop-out and search conditions.

### B. fMRI RESULTS BASED ON VISUAL SEARCH

Through the comparative analysis of imaging data of visual search task, we found that the two search conditions were different in brain imaging. Compared with the search condition, pop-out condition was more active in some brain regions of frontal lobe, temporal lobe, parietal lobe and limbic regions. The activation area of pop-out condition was posterior in the frontal lobe and larger in the temporal and parietal lobe (Table 1).

**TABLE 1.** fMRI activated brain areas in pop-out > search condition.

| Regions | x | y | z | k | T | Z |
| --- | --- | --- | --- | --- | --- | --- |
| Frontal Lobe |  |  |  |  |  |  |
| Right Postcentral gyrus | 60 | -6 | 39 | 68 | 4.4 | 3.73 |
| Right Superior frontal gyrus, medial orbital | 9 | 39 | -9 | 26 | 4.98 | 4.09 |
| Left Superior frontal gyrus, medial orbital | -6 | 48 | -12 | 24 | 3.70 | 3.26 |
| Right Superior frontal gyrus, medial orbital | 6 | 51 | -12 | 26 | 3.41 | 3.05 |
| Left Inferior frontal gyrus, orbital part | -24 | 33 | -9 | 33 | 3.97 | 3.44 |
| Left Inferior frontal gyrus, orbital part | -36 | 39 | -9 | 33 | 3.94 | 3.43 |
| Left Sub-Gyral | -6 | 27 | 0 | 13 | 3.29 | 2.96 |
| Left Middle frontal gyrus | -30 | 18 | 51 | 70 | 3.67 | 3.23 |
| Left Middle frontal gyrus | -24 | 24 | 51 | 70 | 3.52 | 3.13 |
| Temporal Lobe |  |  |  |  |  |  |
| Right Middle temporal gyrus | 57 | 3 | -24 | 83 | 6.56 | 4.92* |
| Right Superior temporal gyrus | 63 | -6 | 6 | 152 | 5.12 | 4.17* |
| Right Middle temporal gyrus | 69 | -30 | 0 | 83 | 4.13 | 3.56 |
| Right Temporal pole: superior temporal gyrus | 39 | 9 | -27 | 44 | 3.82 | 3.34 |
| Left Temporal pole: middle temporal gyrus | -45 | 9 | -27 | 36 | 5.18 | 4.20* |
| Left Superior temporal gyrus | -57 | -18 | 6 | 124 | 5.18 | 4.20* |
| Left Superior temporal gyrus | -54 | -6 | -6 | 292 | 4.66 | 3.89 |
| Left Middle temporal gyrus | -63 | -48 | -6 | 292 | 4.14 | 3.56 |
| Left Middle temporal gyrus | -63 | -12 | -9 | 292 | 4.11 | 3.54 |
| Left Middle temporal gyrus | -54 | -69 | 27 | 70 | 5.34 | 4.29* |
| Left Sub-Gyral | -33 | -54 | 21 | 70 | 3.34 | 3.00 |
| Right Angular gyrus | 54 | -66 | 27 | 64 | 5.28 | 4.26* |
| Right Angular gyrus | 42 | -57 | 27 | 64 | 4.07 | 3.51* |
| Right Middle temporal gyrus | 60 | -60 | 15 | 27 | 4.06 | 3.51* |
| Left Superior temporal gyrus | -42 | -33 | 9 | 15 | 3.15 | 2.85 |
| Parietal Lobe |  |  |  |  |  |  |
| Left Rolandic operculum | -48 | -12 | 15 | 70 | 4.49 | 3.79 |
| Right Postcentral gyrus | 21 | -36 | 72 | 195 | 6.33 | 4.81 |
| Left Precuneus | -6 | -39 | 66 | 256 | 6.3 | 4.79* |
| Left Precuneus | 0 | -63 | 24 | 256 | 4.9 | 4.04* |
| Right Precentral gyrus | 45 | -18 | 60 | 121 | 3.71 | 3.27 |
| Left Angular gyrus | -45 | -63 | 33 | 220 | 5.23 | 4.23* |
| Left Angular gyrus | -42 | -72 | 45 | 220 | 4.87 | 4.02* |
| Left Supramarginal gyrus | -45 | -48 | 24 | 3 | 3.75 | 3.30 |
| Left Postcentral gyrus | -48 | -24 | 60 | 16 | 3.51 | 3.12 |
| Right Rolandic operculum | 51 | -12 | 18 | 223 | 7.11 | 5.17* |
| Limbic Lobe |  |  |  |  |  |  |
| Right Hippocampus | 24 | -9 | -21 | 39 | 6.02 | 4.65* |
| Right Amygdala | 30 | 0 | -24 | 26 | 5.33 | 4.29* |
| Right Parahippocampal gyrus | 36 | -33 | -15 | 19 | 3.82 | 3.34 |
| Left Hippocampus | -21 | -6 | -21 | 41 | 6.38 | 4.83* |
| Left Parahippocampal gyrus | -33 | -3 | -24 | 28 | 4.95 | 4.07* |
| Right Median cingulate and paracingulate gyri | 3 | -36 | 33 | 61 | 5.32 | 4.28* |
| Left Posterior cingulate gyrus | 0 | -45 | 30 | 114 | 5.18 | 4.20* |
| Left Precuneus | -9 | -54 | 12 | 256 | 4.96 | 4.07* |
| Left Fusiform gyrus | -27 | -36 | -15 | 32 | 3.82 | 3.34 |
| Sub-lobar |  |  |  |  |  |  |
| Left Insula | -36 | -15 | 15 | 23 | 4.29 | 3.66 |

Specifically, the frontal lobe included the left middle frontal gyrus, the left inferior frontal gyrus, and the right superior frontal gyrus (medial orbital). The temporal lobe included the right angular gyrus, the right middle temporal gyrus, and the left superior temporal gyrus. The parietal cortex consisted of the bilateral postcentral gyrus and the right rolandic operculum. The limbic regions included the bilateral hippocampus, right medial paracingulate gyri and left fusiform gyrus.

### C. INTEGRATION OF ERP AND fMRI RESULTS

We took the significant activated areas obtained in the fMRI experiment as the source, mapped them to the source model already constructed by ERP data (Figure 7), obtained the corresponding source waveform and scalp distribution, and studied the contribution of these regions to the P300 component. Brain areas that were consistent with ERP and fMRI activation included the right postcentral gyrus of the frontal lobe, the superior frontal gyrus (medial orbital), and the left middle frontal gyrus. The temporal lobes included superior temporal gyrus, middle temporal gyrus, and right angular gyrus. The parietal lobe included the left rolandic operculum, the right postcentral gyrus, the right precentral gyrus, the left angular gyrus, the limbic areas includes the right hippocampus, the left fusiform gyrus, and the sub-lobar area, the left insula (Table 2).

**TABLE 2.** Brain regions with consistent activation of ERP and fMRI.

| Regions | x | y | z | k | T | Z |
| --- | --- | --- | --- | --- | --- | --- |
| Frontal Lobe |  |  |  |  |  |  |
| Right Postcentral gyrus | 60 | -6 | 39 | 68 | 4.4 | 3.73 |
| Right Superior frontal gyrus, medial orbital | 9 | 39 | -9 | 26 | 4.98 | 4.09 |
| Left Superior frontal gyrus, medial orbital | -6 | 48 | -12 | 24 | 3.70 | 3.26 |
| Left Middle frontal gyrus | -24 | 24 | 51 | 70 | 3.52 | 3.13 |
| Temporal Lobe |  |  |  |  |  |  |
| Right Superior temporal gyrus | 63 | -6 | 6 | 152 | 5.12 | 4.17* |
| Right Middle temporal gyrus | 69 | -30 | 0 | 83 | 4.13 | 3.56 |
| Left Superior temporal gyrus | -57 | -18 | 6 | 124 | 5.18 | 4.20* |
| Left Middle temporal gyrus | -63 | -48 | -6 | 292 | 4.14 | 3.56 |
| Right Angular gyrus | 54 | -66 | 27 | 64 | 5.28 | 4.26* |
| Right Middle temporal gyrus | 60 | -60 | 15 | 27 | 4.06 | 3.51* |
| Parietal Lobe |  |  |  |  |  |  |
| Left Rolandic operculum | -48 | -12 | 15 | 70 | 4.49 | 3.79 |
| Right Postcentral gyrus | 21 | -36 | 72 | 195 | 6.33 | 4.81 |
| Right Precentral gyrus | 45 | -18 | 60 | 121 | 3.71 | 3.27 |
| Left Angular gyrus | -45 | -63 | 33 | 220 | 5.23 | 4.23* |
| Limbic Lobe |  |  |  |  |  |  |
| Right Hippocampus | 24 | -9 | -21 | 39 | 6.02 | 4.65* |
| Left Fusiform gyrus | -27 | -36 | -15 | 32 | 3.82 | 3.34 |
| Sub-lobar |  |  |  |  |  |  |
| Left Insula | -36 | -15 | 15 | 23 | 4.29 | 3.66 |

**FIGURE 7.**
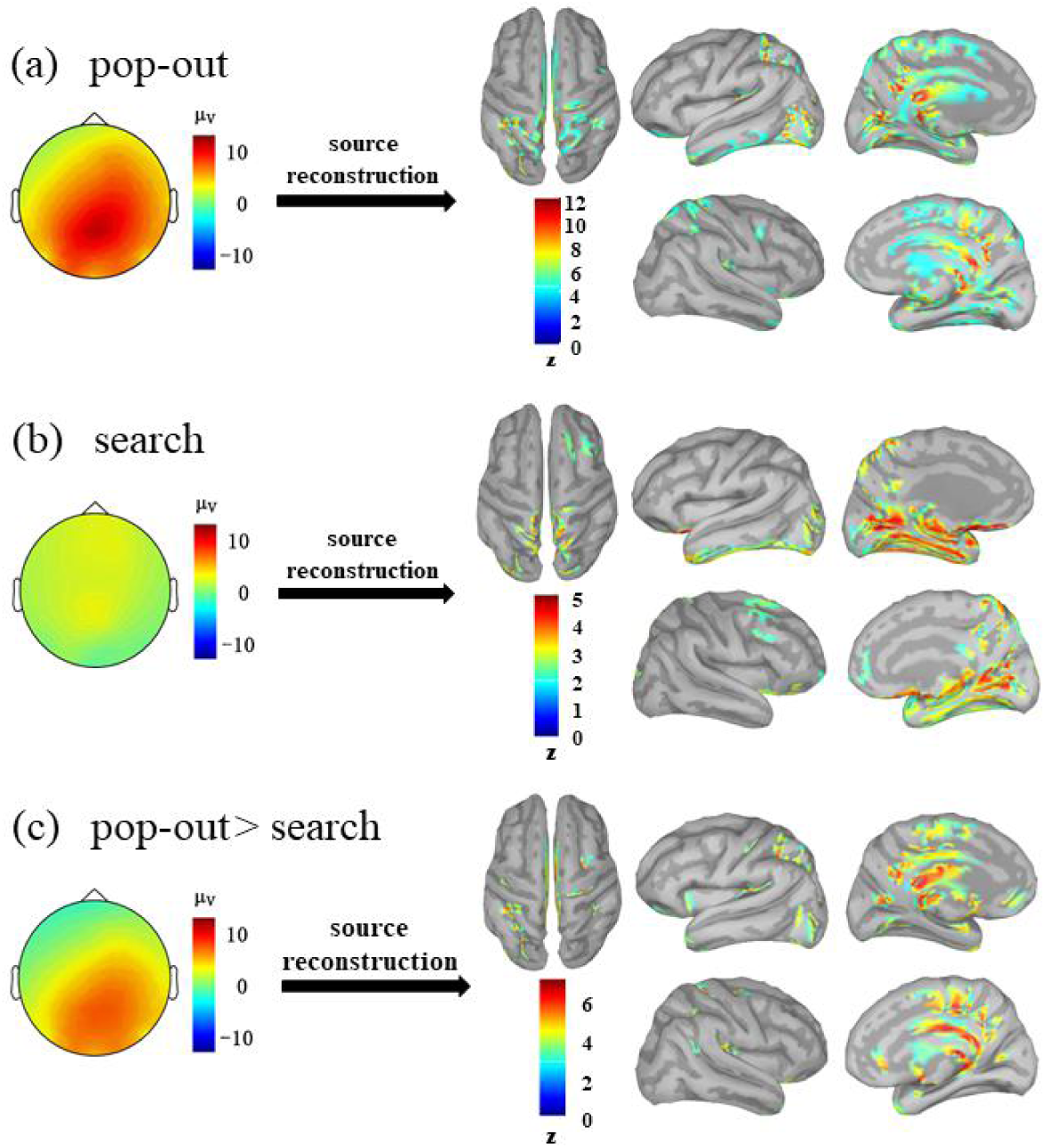
(a) The average source model construction of P300 under the pop-out condition (Active sources with z-score > 4 and adjacent vertexes > 15). (b) The average source model construction of P300 under the search condition (Active sources with z-score > 2 and adjacent vertexes > 15); (c) The average source model construction of P300 under the pop-out > search condition (Active sources with z-score > 3 and adjacent vertexes > 15). From left to right: vertical view of the whole brain (up), lateral view of the left hemisphere (up), medial view of the left hemisphere (up), lateral view of the right hemisphere (bottom), medial view of the right hemisphere (bottom).

The source waveform was extracted by activating the consistent brain region induced by P300 component ERP-fMRI under the two conditions of visual search.

The pop-out condition was found to have left superior frontal gyrus (medial orbital) (t (12) = 4.471, p = 0.001), left angular gyrus (t (12) = 4.61, p = 0.001), and right postcentral gyrus (t (12) = 5.509. p < 0.001) generated significantly larger P300 component than the search condition (Figure 8). The results of ERP-fMRI showed that P300 generated from the left superior frontal gyrus (medial orbital), the left angular gyrus of the parietal lobe, and the right postcentral gyrus.

**FIGURE 8.**
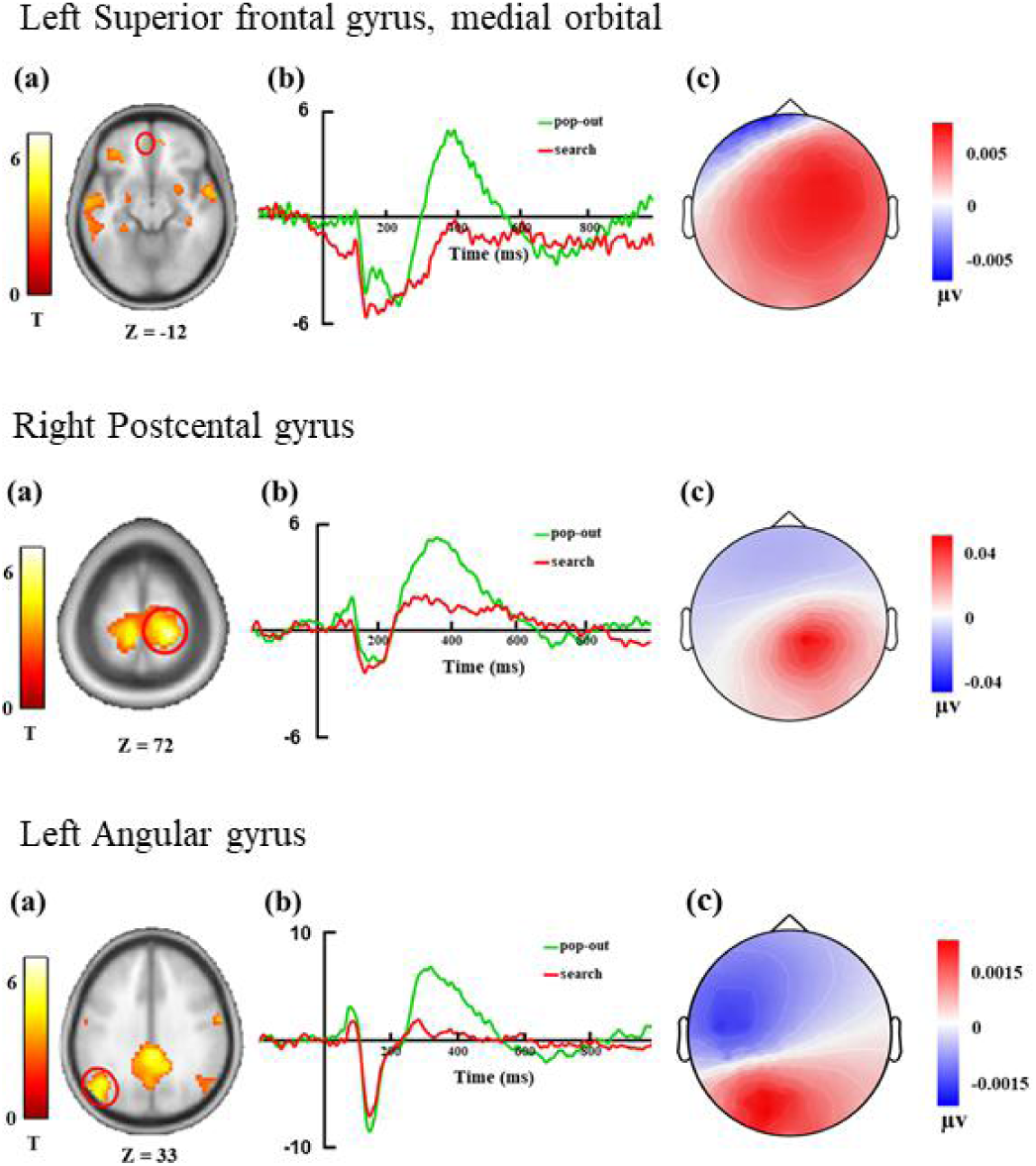
(a) The fMRI indicates significantly activated region under pop > search condition; (b) The waveform of the source activity under the pop > search condition obtained by using the activated region as the source of the brain electrical activity. Green represents pop-out condition and red represents search condition; (c) The scalp mapping of the source activity of the pop > search condition at 350 ms.

## IV. DISCUSSION

### A. ERP DISCUSSIOM

In the ERP experiment, both pop-out and search conditions induced P300, which was distributed in the frontal lobe and parietal lobe. The results confirmed the involvement of the frontal lobe and parietal lobe in attention control [30, 31]. Previous studies have pointed out that the frontal-parietal network is involved in the selective attention of vision [32-34].

The P300 induced by pop-out condition related to bottom-up attention was distributed in the parietal lobe, whereas the P300 induced by search condition related to top-down attention was more distributed in the frontal lobe. Our results also confirmed that in ERP studies on attention in top-down and bottom-up directions, the scalp distribution in the frontal and parietal lobes was different [35-37]. However, other studies have proposed that the frontal-parietal network is a common neural network of bottom-up and top-down attention [38]. In the feature integration theory, feature and object are two basic concepts and individuals conduct parallel processing of features and integrate independent features into objects for serial processing [4]. In an earlier feature integration theory, ERP results showed that the parallel feature-present condition induces larger P300 in the parietal lobe than serial feature-absent condition [39, 40]. This forms an important basis for evaluating the visual search of parallel and serial model study, with visual search “feature-present” and “feature-absent” condition respectively corresponding to the “parallel” and “serial” processing. The difference between our pop-out and search tasks lies in the presence and absence of the “color” feature of the target and the distractor. Our results also confirmed that the pop-out task with the “color” feature induces a larger P300 in the parietal lobe than the search task without the “color” feature.

Although both pop-out and search induced P300, pop-out induced larger P300. Arguably, P300 reflects the process of cognitive processing of information and it’s amplitude reflects the attention resources assigned to tasks by individuals [26]. The perceptual load theory suggested that the level of perceptual load determines the allocation of resources in the process of selective attention [41]. In the pop-out condition, the target is more recognizable, and the individual allocates more attention resources to the target than to the distractor, thus inducing a larger P300. According to the similarity theory of visual search, the higher the similarity between target and distractor, the more difficult the visual search will be [6]. The similarity between the target and distractor in the search condition was higher than that in the pop-out condition, resulting in a higher level of memory load for the search condition. Our results showed that different memory loads induce different P300 amplitudes, and P300 amplitudes at high load was lower than those at low memory load [42, 43].

### B. fMRI DISCUSSIOM

In the study, pop-out condition showed more activation in the frontal lobe, temporal lobe, parietal lobe and some limbic areas than the search condition. In some fMRI studies using the visual oddball paradigm, researchers have found that the brain regions involved in the activation of the target detection process include the inferior parietal lobule and postcentral sulcus of the parietal lobe, and the medial frontal gyrus, medial orbitofrontal cortex and superior frontal gyrus [44-46]. Our results supported that the detection of targets in visual search involves a wide range of brain regions [47].

The pop-out task involves stimulus-driven attention control, whereas the search task involves goal-driven attention control. It can thus, be speculated that different attentional control is associated with activations in different brain areas. Previous studies have pointed out that individuals performing different attention tasks experience activations corresponding to different brain networks. For example, the temporal-parietal cortex and inferior frontal gyrus participate in stimulus-driven attention, the medial parietal lobe and superior frontal gyrus participate in top-down attention of target control [1]. We found that the pop-out task controlled by bottom-up attention corresponded with more activation in the temporal lobe, parietal lobe and posterior frontal lobe, which was consistent with the earlier results.

In the visual search task, subjects were required to remember the characteristics of the target, and then to search and respond to the stimuli containing the target and the distractor, which involves both attention and working memory. Attention is closely related to working memory. By paying attention to representation and early perception, the target that need to be remembered are encoded and stored, and then the memory content is extracted to guide the attention to the stimuli matching the target in the field of vision [48-50]. A meta-analysis of fMRI studies on working memory of language indicate that working memory involves activity in brain regions such as frontal lobe, parietal lobe, cerebellum, and fusiform gyrus of limbic system [51]. In fact, the studies on visual working memory have implicated the activities of the inferior frontal gyrus and the middle frontal gyrus participate in working memory [52, 53]. In our study, the pop-out task showed strong activity in the middle frontal gyrus, inferior frontal gyrus, parietal lobe and fusiform gyrus of the frontal lobe, and thus, our results reflected the neural overlap between visual search attention and working memory.

### C. INTEGARTION OF ERP fMRI DISCUSSIOM

We located the source of the P300 components induced by pop-out and search through the integration of ERP and fMRI data, and found that P300 generated from the left superior frontal gyrus (medial orbital), the left angular gyrus, and the right postcentral gyrus. The pop-out task was associated with stimulus-driven bottom-up attention, in which the target stimulus became prominent and automatically attracted the participants’ attention. In the present study, the pop-out induced P300 was distributed in the parietal lobe, the left angular gyrus, and right postcentral gyrus, with the latter having stronger activation. This indicates that stimulus-driven bottom-up attention may be related to the activities of the angular gyrus and postcentral gyrus.

Moreover, the visual search task involves the parietal lobe and the frontal lobe. In one study, P300 source analysis was performed on the target and distractor responses of vision, and it was found that the brain regions involved in the processing of the target and non-target were different. The processing of the target was driven by the stimulus, and the P300 component was linked to the parietal lobe and the inferior temporal gyrus, rather than the processing of target P300 involving the insula and cingulate [54]. In an oddball paradigm with words as stimulus, the researchers found that the average P300 of individual subjects and groups was mainly distributed in the center of the parietal lobe, and the peak of P300 was distributed in the left parietal lobe. Especially, P300 generated from the intraparietal sulcus and the superior parietal lobule [55]. In another study with cerebral infarction patients as the subjects, the P300 amplitude of lesions in the superior temporal gyrus, inferior frontal gyrus and prefrontal area of the patients was significantly reduced, which led to the conclusion that the temporal lobe and frontal lobe are the important brain regions for P300 production [56]. Consistently, our findings confirmed the contribution of the frontal and parietal lobes to the P300 component.

In addition, a number of studies have found that the temporal lobe is also an important region of P300 [57, 58]. Review studies on the source of P300 indicated that P300 nerve electrical events generated from the temporoparietal junction, medial temporal lobe, and lateral frontal lobe [59]. In our experiment, although two visual search tasks were found to be activated in the superior and middle temporal gyrus, P300 component did not show significant activation in these areas. Interestingly, we found the angular gyrus to be the source of P300. Since the angular gyrus is located at the tail end of the superior temporal sulcus, it is possible that the temporal lobe region to some extent, is also involved in the production of P300.

Currently, there are many hypotheses about the source of P300, and the most recognized one is the multi-source hypothesis. This hypothesis suggests that P300 source from cortical and subcortical structures, including multiple brain regions such as frontal lobe, parietal lobe, temporal lobe, and limbic system [60]. Based on ERP and fMRI data, we found that P300 is associated with the activation of these brain regions, which also supported the multi-source hypothesis. In our study, the source of P300 was further narrowed down to defined brain region, including the left superior frontal gyrus (medial orbital), the left angular gyrus, and the right posterior central gyrus.

Furthermore, EEG signal and fMRI data were collected separately. Although the experimental tasks of the two methods were the same, the subjects may have inconsistent cognitive neural processes, which may affect the results. A growing number of studies are using simultaneous EEG and fMRI data to ensure that both types of data reflect the same neural processes and that the subjects use the same strategies for both types of data. Generally, P300 is an important marker of the cognitive process, and the research on the source of P300 is crucial in furthering the understanding of its cognitive significance and clinical application. Studies have found that P300 induced by patients with dementia [61], cerebral infarction [62], and Parkinson’s disease [63] were different from that of healthy individuals. ERP-fMRI technology can be used to increase the research on the source of patients’ P300, which will provide a basis for medical researchers to develop treatment plans and help patients to overcome their cognitive deficits.

## V. CONCLUSION

We induced P300 by visual search task and found that P300 dominated by different attention process was different. P300 induced by bottom-up attention (pop-out) is more concentrated in the parietal lobe, whereas P300 induced by top-down attention (search) is more distributed in the frontal lobe. Using ERP-fMRI integration analysis, left angular gyrus, right postcentral gyrus of and the left superior frontal gyrus (medial orbital) were the neural source. Our finding suggested that P300 source mainly from the frontal-parietal lobe, indicating that the frontal-parietal lobe plays a key role in the process of attention.

